# Sex makes them sleepy: host reproductive status induces diapause in a parasitoid population experiencing harsh winters

**DOI:** 10.1101/371385

**Authors:** K. Tougeron, J. Brodeur, J. van Baaren, D. Renault, C. Le Lann

## Abstract

When organisms coevolve, any change in one species can induce phenotypic changes in traits and ecology of the other species. The role such interactions play in ecosystems is central, but their mechanistic bases remain underexplored. Upper trophic level species have to synchronize their life-cycle to both abiotic conditions and to lower trophic level species’ phenology and phenotypic variations. We tested the effect of host seasonal strategy on parasitoid diapause induction by using a holocyclic clone of the pea aphid *Acyrthosiphon pisum* producing asexual and sexual morphs that are viviparous females (i.e. laying embryos) and oviparous females (laying eggs), respectively, the latter being only present at the end of the growing season. *Aphidius ervi* parasitoids from populations of contrasted climatic origin (harsh vs. mild winter areas) were allowed to parasitize each morph in a split-brood design and developing parasitoids were next reared under either fall-like or summer-like temperature-photoperiod conditions. We next examined aspects of the host physiological state by comparing the relative proportion of forty-seven metabolites and lipid reserves in both morphs produced under the same conditions. We found that oviparous morphs are cues *per se* for diapause induction; parasitoids entered diapause at higher levels when developing in oviparous hosts (19.4 ± 3.0%) than in viviparous ones (3.6 ± 1.3%), under summer-like conditions (i.e., when oviparous aphids appear in the fields). This pattern was only observed in parasitoids from the harsh winter area since low diapause levels were observed in the other population, suggesting local adaptations to overwintering cues. Metabolomics analyses show parasitoids’ response to be mainly influenced by the host’s physiology, with higher proportion of polyols and sugars, and more fat reserves being found in oviparous morphs. Host quality thus varies across the seasons and represents one of the multiple environmental parameters affecting parasitoid diapause. Our results underline strong coevolutionary processes between hosts and parasitoids in their area of origin, likely leading to phenological synchronization, and we point out the importance of such bottom-up effects for trait expression, and for the provision of ecosystem services such as biological control in the context of climate change.

## Introduction

Interacting individuals from two biological entities can adjust their phenotypes in response to cues from each other, even when these cues vary over time (Agrawal 2001). Beneficial or antagonistic interactions, from mutualism to parasitism, predation and competition may lead to adaptive phenotypic responses. When interactions persist over generations, coevolution can occur and species adapt to the interacting species’ life history traits, phenology and ecology (Agrawal 2001, Ellers et al. 2012). Interaction norms (Thompson 1988) arise from ecological responses of interacting organisms in varying environments, as any phenotypic change occurring in one “partner” species can cascade to the other species’ phenotype (Fordyce 2006, Hughes 2012). Cues produced by one interacting species may indirectly inform the other species of environmental changes. For example, plant senescence in fall can inform herbivorous insects of upcoming detrimental winter conditions and induces phenotypic changes (e.g. diapause induction) or migration behaviour (Archetti et al. 2009).

Parasitoids are excellent models to study phenotypic expression in interacting species because they are strongly influenced during immature stages by changes in nutritional and physiological quality of their host (Godfray 1994). Diapause is an important ecological process in insects allowing them to survive recurrent unfavorable environmental conditions (Tauber et al. 1986). For parasitoids, diapause also contributes to maintaining synchronization with their host’s seasonal reproductive cycle; it is induced before suitable hosts vanish from the environment (Lalonde 2004). As in most insects, diapause in parasitoids is mainly induced by abiotic cues perceived either by the generation that will enter diapause, or by the maternal generation (Tauber et al. 1986). A few studies also reported that diapause in parasitoids can be triggered by the onset of host diapause (Polgár and Hardie 2000, Gerling et al. 2009), or through intraspecific competition for hosts (Tougeron et al. 2017a). However, whether the phenotype of a non-diapausing host can influence parasitoid diapause remains poorly studied.

Aphids are hosts for Aphidiinae parasitoids and can have very complex cycles showing seasonal alternation between morphs with asexual and sexual reproduction (Dixon 1985). Asexual females reproduce parthenogenetically and lay live offspring (i.e. viviparity) whereas sexually reproducing females produce eggs (i.e. oviparity) after mating with males. Sexual aphid morphs are present at higher proportions in harsh than in mild winter climates (Dedryver et al. 2001), and they represent the last hosts available for aphid parasitoids before winter as they produce overwintering eggs in fall (Leather 1992). Consequently, sexual morphs have been suggested to promote diapause in parasitoids, indicating a host physiological effect (Polgár et al. 1991, 1995, Christiansen-Weniger and Hardie 1997). No mechanistic understanding of this phenomenon has been proposed and the effects of the host morph have not been disentangled from confounding factors such as host genotype and geographic origin, host size, abiotic conditions, or the season at which hosts are sampled in the fields. Hosts and parasitoids have coevolved over long periods of time, they respond to similar seasonal cues and the physiological syndrome associated with overwintering is highly conserved among insects (Tauber et al. 1986, Denlinger 2002). As a result, the related physiological state of the host may represent a reliable signal of upcoming seasonal changes for parasitoids.

Hormones, fats, carbohydrates and other types of metabolites are involved in the regulation of overwintering and diapause expression in insects (Chippendale 1977, Christiansen-Weniger and Hardie 1999, Denlinger 2002, Sinclair and Marshall 2018). In aphid parasitoids, metabolomic and proteomic profiles differ between diapausing and non-diapausing individuals, with higher amounts of sugars, polyols and heat shock proteins being found in diapausing parasitoids (Colinet et al. 2012). In aphids, morphs differ in morphology and physiology; oviparous females accumulate reserves to produce energetically costly diapausing eggs (Le Trionnaire et al. 2008) with cryoprotectant compounds such as mannitol and glycerol (Sömme 1969), whereas viviparous females metabolize energetic resources rapidly to produce embryos. Aphids’ triglyceride reserves change quantitatively and qualitatively across the seasons with alternating morphs (Greenway et al. 1974). Immature parasitoids are known to consume sugars and lipids from their hosts (Jervis et al. 2008) and are therefore influenced by host reserves for their growth and development.

We questioned the extent to which oviparous and viviparous morphs of a single clone of the pea aphid *Acyrthosiphon pisum* (Harris) (Hemiptera: Aphididae) influences winter diapause expression in the parasitoid *Aphidius ervi* Haliday (Hymenoptera: Braconidae) under summer and fall conditions. Under laboratory conditions and using a split-brood design, we compared the response to two aphid morphs of two populations of parasitoids from mild (France) and harsh (Canada) winter areas that differed in their level of diapause expression (Tougeron et al. 2018). In *Aphidius* species, winter diapause is initiated at the prepupal stage within the aphid mummy (i.e. dead aphid containing a developing parasitoid) following stimuli perceived by the mother or early developmental stages (Brodeur and McNeil 1989, Tougeron et al. 2017b). We hypothesized that parasitoids of both populations developing in oviparous hosts enter diapause at higher proportions than those developing in viviparous hosts, independently of photoperiod and temperature. We predicted this pattern to originate from differences in aphids’ physiological contents. We thus performed physiological analyzes to measure lipid content and quantify aphid morphs metabolites. We also hypothesized parasitoids from the mild winter area to be less responsive to diapause-inducing cues from the host and the environment, because parasitoid populations should be adapted to climatic conditions and to the relative occurrence of sexual hosts in their respective areas of origin.

## Material and Methods

### Biological materials

Two populations of the parasitoid *A. ervi* were collected in 2015 at the mummy stage in pea fields from two contrasted climatic origins: near Montréal, QC, Canada (45.584°N, 73.243°W; harsh winter area) and near Rennes, France (48.113°N, 1.674°W; mild winter area). One population per geographic origin was used as high gene flow has been reported in *A. ervi* populations, which therefore present little genetic differentiation (Hufbauer et al. 2004). Even if gene flow was weak, we would expect higher differences between Canadian and French populations than among populations of a same location. Parasitoids were then reared under controlled conditions using a cyclically parthenogenetic clone (clone F2-X9-47) of the pea aphid *A. pisum* provided by INRA Le Rheu, France, and known to produce both oviparous and viviparous aphid morphs (Jaquiéry et al. 2014). The symbiotic load of the aphid clone we used was not assessed, but symbionts present in the grandparent generation from which our clone comes from had been identified. Half of the grandparent generation was associated with *Serratia symbiotica*, the other half had no secondary endosymbionts (J. Jaquiéry pers. comm.). It is thus likely that our clone was inhabited by *S. symbiotica*. All insects were maintained on fava beans *Vicia faba* (Fabaceae) at 20 °C, 70% relative humidity (RH) and 16:8 h Light:Dark (L:D) photoregime.

### Production of sexual and asexual hosts

Three aphid morphs were used in the experiments; oviparous females (O), viviparous females (V) and a control treatment for viviparous females (C), as detailed below.

Three parthenogenetic *A. pisum* adult females from the aphid culture were put on bean plants (N=15) and allowed to lay larvae during four days at 20 °C, 70% RH, 16:8 h L:D. Females were then removed and infested plants were put in a growing chamber at 17 °C, 70% RH, 12:12 h (L:D), and under 36W, IRC 85, 6500 K day-light type fluorescent tubes to induce the production of sexual aphids (Le Trionnaire et al. 2009). At each generation, plants were renewed, and less than five aphids were maintained per plant to prevent formation of winged individuals due to overcrowding (Hardie 1980). As embryos directly detect photoperiodic cue through the cuticle of the grand-mother (Le Trionnaire et al. 2008), the first sexual aphids: males (∼20%) and oviparous females (30 to 60%) were formed, along with asexual aphids (20 to 50%): sexuparous (a particular type of parthenogenetic females producing sexual morphs) and viviparous aphids (parthenogenetic females producing only parthenogenetic morphs), after three generations under these conditions. As sexuparous and viviparous aphids cannot be distinguished morphologically, they were indistinctly considered as the “viviparous female” treatment. However, a control group of viviparous parthenogenetic females (C) was produced by rearing aphids under non-sexual-inductive conditions (20 °C, 70% RH, 16:8 h L:D). This treatment controls for potential stress effects of the sexual-inductive conditions on the aphid, and allows to solely measure the response of viviparous aphids as sexuparous are not produced under this condition (Dixon 1985). Oviparous aphid morphs were differentiated from viviparous ones under a stereo microscope (x10) by observing the morphology of their legs: oviparous female aphids have rhinaria on the tibia, and have a femur of the same width as the tibia, and viviparous females have a wider tibia than the femur without rhinaria (Lamb and Pointing 1972, Hullé et al. 2006). Aphid males were not included in our analyses since *A. ervi* does not parasitize them, probably because they are too small and have lower energetic reserves than female morphs (Tougeron et al., unpublished data).

### Diapause induction

Aphid mummies from the colonies were isolated in a small gelatin capsule until parasitoid emergence. Newly emerged parasitoids were put in a 5 cm plastic tube for mating (5 females with 2 males) for 24 h, and were fed with a 70% diluted honey solution. Maternal genotype, egg-laying order in different aphid morphs, in addition to parasitoids’ age or host preference may affect diapause induction (Brodeur and McNeil 1989). To consider these potential effects, twelve *A. ervi* females were individually allowed to parasitize 16 adult aphids of the same age and size within the same cohort and of each of the three morph types (oviparous female, viviparous female, control viviparous females produced under non-sexual-inductive conditions, N=48 aphids offered for parasitism per female wasp) for 12 h over three consecutive days, by alternating the order of presentation of aphid morphs among females. Parasitoids rested at night, with an access to diluted honey. Aphids were introduced in a plastic tube (10 × 3 cm) and were given a few minutes to settle on a bean cut plant, after which a parasitoid was introduced into the tube. Four parasitoid females were first individually put in presence of oviparous aphids, then moved to a second tube with control viviparous aphids and next moved to a third tube containing viviparous aphids (OCV). Four other females were first offered viviparous aphids (VOC), and the last four females were first offered control viviparous aphids (CVO) (Fig. 1).

**Figure 1:**
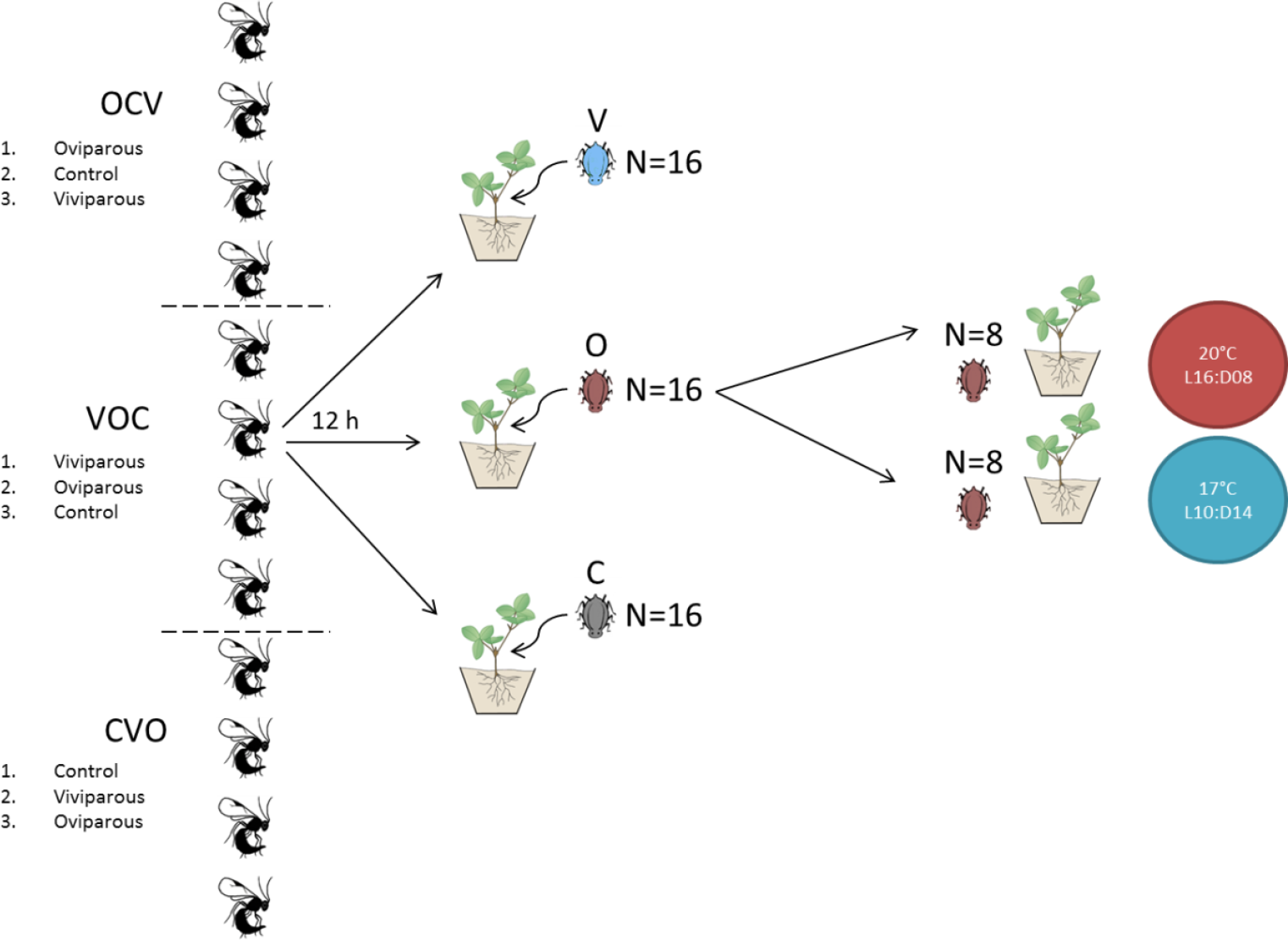
Experimental design for diapause induction in the parasitoid *Aphidius ervi*. Twelve parasitoid females were individually allowed to parasitize 16 *Acyrthosiphon pisum* from each of the three host morphs for 12 h: oviparous (O), viviparous (V) and viviparous control (C). First contact (parasitism sequence) with an aphid was alternated between the three morphs (OCV, VOC, CVO). Following parasitism, the aphid cohort was split in two and individuals were reared under a diapause-inductive condition (17 °C 10:14 h L:D) or a non-diapause-inductive condition (20 °C 16:8 h L:D). This protocol was repeated twice for parasitoid populations originating from mild or harsh winter.

After each oviposition period, the 16 potentially parasitized aphids of each morph type were transferred by group of 8 on two bean plants. Plants were next enclosed into micro-perfored plastic bags and placed at either 20 °C, 16:8 h (L:D) (summer-like conditions not inducing diapause in *A. ervi*) or 17 °C, 10:14 h (L:D) (autumn-like conditions inducing diapause) (Tougeron et al. 2017b). When the plants began to wilt, aphids were transferred to another plant with a small paintbrush. Mummification was checked daily and newly-formed mummies were placed individually into gelatin capsules, and remained under their respective temperature and photoperiod treatments until adult emergence. Mummies from which no parasitoid had emerged 15 days after mummification were dissected, and the content was recorded as dead parasitoids or diapausing individuals (golden-yellow prepupae, Tougeron et al. 2017b). This experiment was repeated twice per parasitoid population; diapause levels were thus calculated among the offspring of 24 females for each treatment. Patterns were consistent in each of the repeated experiments. Our split-brood family design also allowed comparing reaction norms (RN) of diapause levels in the offspring of each parasitoid female from each population, both within morphs at different abiotic conditions, and within abiotic conditions among morphs. We have excluded “control” morphs from the RN analysis as their effect on diapause induction did not differ from viviparous morphs.

The aphid morph (individual differences within a population due to developmental plasticity) and the aphid clone (differences in reproduction modes genetically determined between populations) may both influence parasitoid diapause. To consider this aspect, we compared the incidence of diapause when parasitoids developed in the cyclically parthenogenetic clone (holocyclic, i.e., alternating between sexual and sexual morphs) described above and in an obligate parthenogenetic clone, producing only viviparous females (anholocyclic clone F2-X9-19; Jaquiéry et al. 2014). To achieve this goal, five *A. ervi* females were individually allowed to sequentially parasitize 35 viviparous aphids of each clone during 12 h. Parasitized hosts were next placed at 17 °C 10:14 h (L:D), and diapause induction was measured as described above. We excluded any clone effect because diapause incidence was similar for parasitoids developing in viviparous aphids of either the holocyclic (59.9 ± 10.1%, n=132 mummies) or the anholocyclic (66.0 ± 7.7%, n=112 mummies) clone (GLM, p=0.97). The cyclically parthenogenetic clone was thus used for the experiments.

### Metabolomic analyses and lipid reserves

As sexual morphs could only be produced at 17°C, we compared non-parasitized apterous adult aphids of viviparous and oviparous females of the same age (between 24 and 48 h after imago molt), produced under the same conditions used for the diapause experiment (at 17 °C, 12:12 h (L:D)). Samples were kept at −20 °C for metabolomic and lipid analyses. They were dried out for 2 days in a freeze-dryer and their dry mass measured using a Mettler-Toledo precision scale (accurate to 0.001 mg). Viviparous aphids’ dry mass ranged from 0.280 mg to 0.742 mg, and oviparous aphids’ dry mass ranged from 0.358 mg to 0.739 mg.

For metabolic analyses, 18 aphids of each morph (viviparous and oviparous females) were used. Nine replicates were analyzed for each morph condition, each consisting of a pool of two aphid females. The samples were put in 600 µL of chloroform-methanol (1:2) solution and homogenized using a tungsten-bead beating apparatus at 30 Hz for 1.5 min. Then, 400 µL of ultrapure water was added to each tube and samples were centrifuged at 4 °C, 4,000 g for 5 min. Finally, 90 µL of the upper aqueous phase containing metabolites were transferred to chromatographic vials. Injection order of the samples was randomized prior mass spectrometry detection. Metabolomic fingerprinting process was performed following the protocol of Khodayari et al. (2013). Chromatograms were analyzed using XCalibur software (Thermo Fischer Scientific, Waltham, MA, USA). We accurately quantified 47 metabolites: 14 amino acids, 11 sugars / sugar phosphates, 8 organic acids, 7 polyols, 4 other metabolites and 3 amines (Table 1).

**Table 1:** Metabolites detected in each of the two morphs (viviparous and oviparous females) of the pea aphid, *Acyrthosiphon pisum*. Each metabolite has been found in each morph. Abbreviations used on Figure 3 are in brackets.

| Amino acids | Organic acids |
| --- | --- |
| Alanine (Ala) | Citric acid (Cit_Ac) |
| Aspartic acid (Asp_Ac) | Galacturonic acid (Gal_Ac) |
| Citrulline (Citr) | Glyceric acid (Glyc_Ac) |
| Glutamic acid (Glu) | Lactic acid (Lact_Ac) |
| Glycine (Gly) | Malic acid (Mal_Ac) |
| Isoleucine (Ile) | Phosphoric acid (Phos_Ac) |
| Leucine (Leu) | Pipecolic acid (Pipe_Ac) |
| Lysine (Lys) | Quinic acid (Quin_Ac) |
| Ornithine (Orn) |  |
| Proline (Pro) | **Sugars and sugar phosphates** |
| Serine (Ser) | Arabinose |
| Valine (Val) | Fructose |
| Threonine (Thr) | Fructose-6-phosphate (F6P) |
| Phenylalanine (Phe) | Galactose |
|  | Glucose |
| **Polyols** | Glucose-6-phosphate (G6P) |
| Adonitol | Maltose |
| Arabitol | Mannose |
| Galacticol | Ribose |
| Glycerol | Saccharose |
| Inositol | Trehalose |
| Mannitol |  |
| Xylitol | **Other metabolites** |
| **Amines** | Gluconolactone (GNL) |
| Cadaverine (Cad) | Gamma aminobutyric acid (GABA) |
| Triethanolamine (TEA) | Glycerol-3-phosphate (Gly3P) |
| Putrescine (Put) | Dopamine (Dop) |

Lipid contents were measured using 52 oviparous females and 23 viviparous females. Each dry aphid was left for two weeks in a microtube containing 1 mL of chloroform-methanol solution (2:1) to extract lipids (Terblanche et al. 2004). Aphids were then rinsed with the same solution, and placed back in the freeze-dryer for 24 h to eliminate the residues of the extracting solution and next weighted again to measure fat content (= fat mass (mg) / lean dry mass (mg), Colinet et al. 2007).

### Statistical analyses

Generalized linear mixed-effects models (GLMM) with binomial distributions were fit to the data using the *lme4* package. The response variable was the proportion of diapausing parasitoids; the origin of the parasitoid population (Canada vs. France), the host morph (three modalities, O, V, C), the temperature/photoperiod conditions (17°C 10:14h vs 20°C 16:8h), and their interaction, were considered as fixed factors; the identity of each parasitoid female and the egg-laying (parasitism) order were considered as random effect factors in the models. As diapause incidence differed between parasitoid populations (GLMM, χ^2^=216, df=1, p<0.001), data from both populations were analyzed separately using similar GLMMs. Significance of each term in the model was analyzed using the package *car*.

For metabolite data, concentrations of the compounds were first log-transformed. Then, a Principal Component Analysis (PCA) was performed to detect which metabolites (expressed in nmol.mg^-1^) differed the most between host morphs. Log-transformed metabolite concentrations were then summed up within each category (Table 1) and another PCA was performed using metabolite groups as discriminatory factors. An ANOVA with FDR-adjusted p-values was next performed to compare concentrations of each metabolite between morphs. Finally, an ANOVA tested differences in fat content between oviparous and viviparous morphs. All statistical analyses were carried out using the R software (R Core Team 2017).

## Results

### Diapause incidence in the parasitoid A. ervi

In the Canadian (harsh winter area) population, diapause levels were affected by host morph (GLMM, χ^2^=12.6, df=2, p<0.001; Fig. 2) and abiotic conditions (GLMM, χ^2^=250.0, df=1, p<0.001), with an interaction effect as host morphs influenced parasitoid diapause incidence only at 20 °C 16:8 h (L:D) (GLMM, χ^2^=16.9, df=2, p<0.001). Diapause incidence was higher at 17 °C 10:14 h L:D than at 20 °C, 16:8 h L:D, for the Canadian population (76.9 ± 2.5% *vs.* 9.0 ± 1.5%, respectively). At 20 °C, 16:8 h L:D, diapause incidence was higher when Canadian parasitoids developed in oviparous aphids (19.4 ± 3.0% s.e.) than in viviparous aphids (3.6 ± 1.3%, z=-4.3, p<0.001) or viviparous control aphids (3.8 ± 1.4%, z=-3.9, p<0.001).

**Figure 2:**
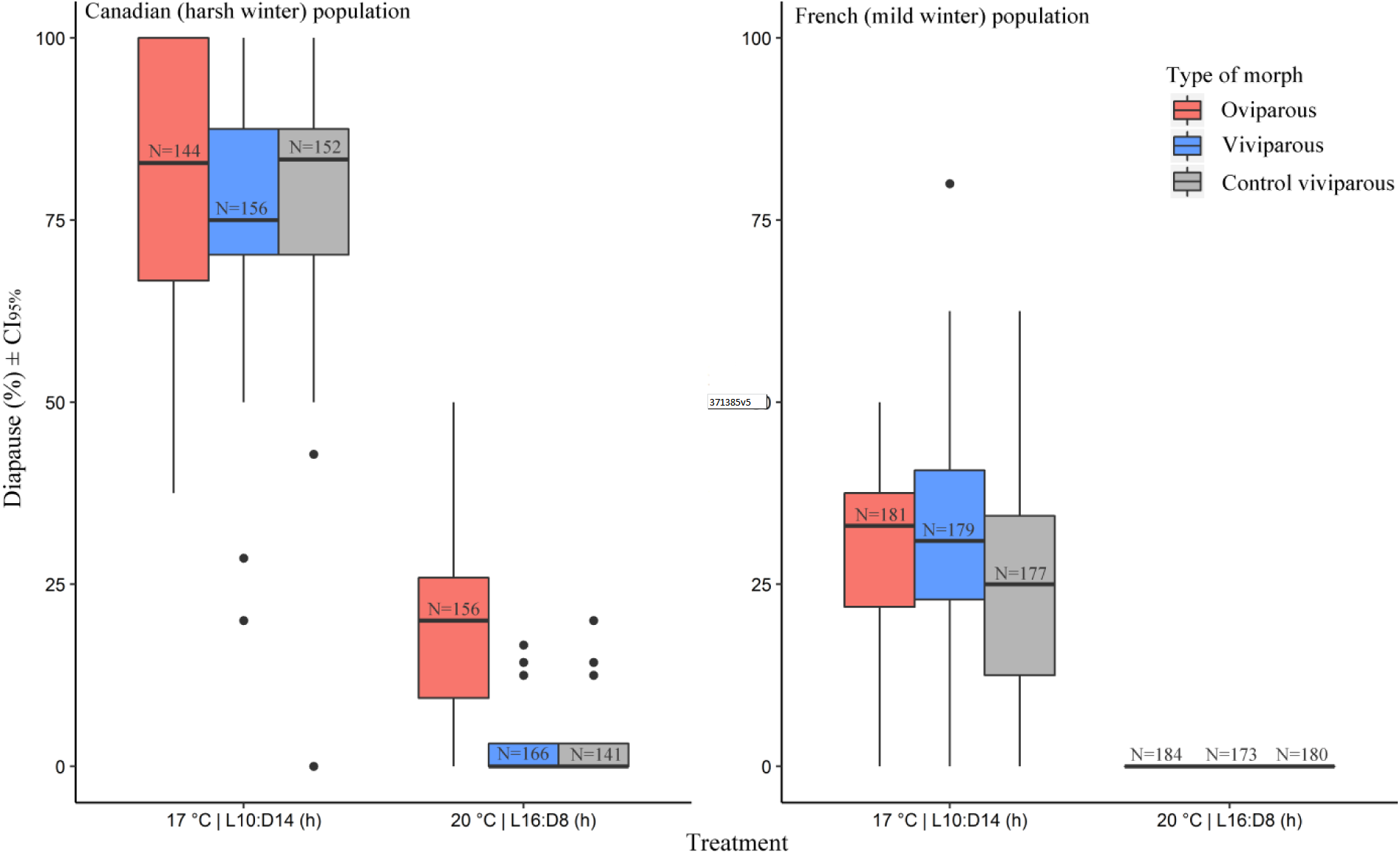
Percent diapause incidence (± CI_95%_) in two *Aphidius ervi* populations. *Left:* Canadian population naturally experiencing harsh winter. *Right*: French population naturally experiencing mild winter. For both populations, three different morphs of the pea aphid *Acyrthosiphon pisum* (oviparous sexual females, viviparous parthenogenetic females produced under sexual-inductive conditions, and control viviparous females produced under non-sexual-inductive conditions) were used for parasitoid development, under two abiotic conditions (17 °C, 10:14 h L:D or 20°C, 16:8 h L:D). For each treatment, N represents the total number of parasitoid mummies used to calculate diapause incidence.

In the French (mild winter area) population, the host morph did not influence parasitoid diapause (GLMM, χ^2^=1.84, df=2, p=0.39), abiotic conditions did influence parasitoid diapause (GLMM, χ^2^=237.9, df=1, p<0.001), but no interaction effect can be interpreted since no diapause was expressed for the French population at 20 °C, 16:8 h L:D. Diapause incidence was higher at 17 °C 10:14 h L:D than at 20 °C, 16:8 h L:D, for the French population (27.9 ± 2.1% *vs.* 0%, respectively). Random factors female identity and host exposition order had negligible effects on total variance explained in both our models for both populations (variance ≤0.02).

Some female parasitoids produced offspring that had stronger responses to changes in host morph or abiotic conditions than offspring of other females (Fig. 3). Data for each female are made available as a supplementary material sheet. In some broods, there was no variation in diapause plasticity in response to different biotic (morphs) or abiotic (photoperiod and temperature) conditions (RN slope = 0). In the Canadian population at 17°C 10:14 h L:D, reaction norm slopes (i.e., diapause level variations between conditions within a single brood) ranged from −71% to 48%, for the diapause response to either oviparous or viviparous morphs Fig. 3A). At 20°C 16:8 h L:D, these RN slopes ranged from −50% to 12% (Fig. 3B). In the French population at 17°C 10:14 h L:D, RN slopes ranged from −29% to 38% for the diapause response to either oviparous or viviparous morphs (Fig. 3C).

**Figure 3:**
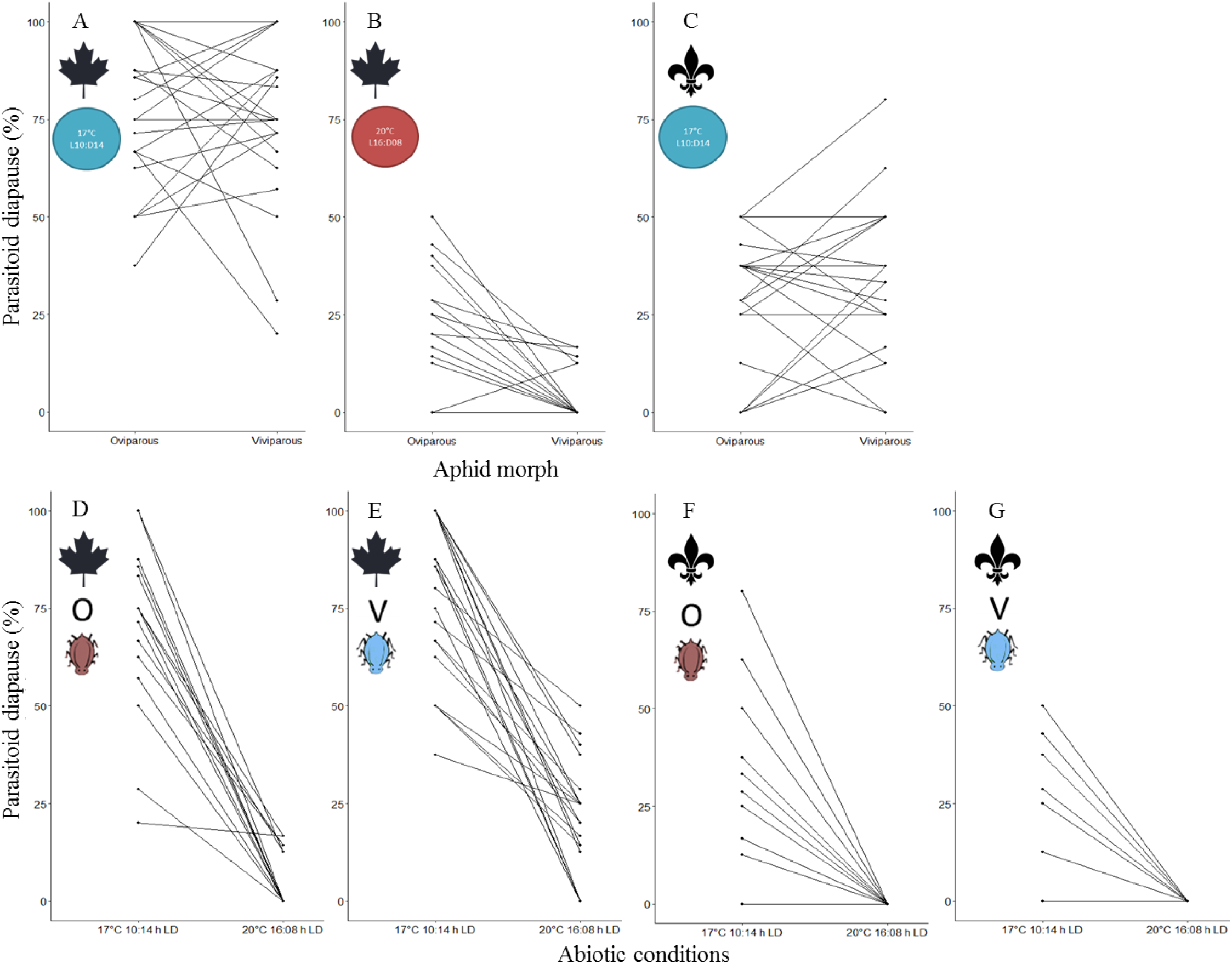
Reaction norms (RN) of diapause levels in the offspring of each parasitoid female from each parasitoid population (Canadian: 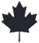 and French: 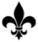), both within morphs at different abiotic conditions (top panel, **A & C:** 17°C 10:14 h L:D, **B:** 20°C 16:8 h L:D)), and within abiotic conditions between morphs (bottom panel, **D & F**: oviparous morphs, **E & G**: viviparous morphs). RN for the French population at 20°C 16:8 h L:D are not displayed as no diapause was observed under these conditions. N=24 parasitoid female per condition. Note that some lines may be overlapping.

In the Canadian population, for parasitoids developing in viviparous morphs, RN slopes ranged from −100% to −3% (Fig. 3D), and for parasitoids developing in oviparous morphs, RN slopes ranged from −100% to −12% (Fig. 3E), for the diapause response to abiotic conditions (17°C 10:14 h L:D *vs.* 20°C 16:8 h L:D). In the French population, for parasitoids developing in viviparous morphs, RN slopes ranged from - 80% to 0% (Fig. 3F), and for parasitoids developing in oviparous morphs, RN slopes ranged from −50% to 0% (Fig. 3G) for the diapause response to either abiotic conditions.

### Metabolomic analyses and lipid reserves of aphid host morphs

All measured compounds were found in both aphid morphs. The first and second principal component (PC1 and PC2, respectively) of the PCA, accounted for 37.1% and 26% of the total inertia, respectively (Fig. 3). Oviparous and viviparous female hosts were separated on PC1, with oviparous females exhibiting significantly higher concentrations of trehalose, ribose, arabitol, gamma aminobutyric acid and mannose than viviparous ones (ANOVA, df=1, p<0.05) (Fig. S1). Conversely, viviparous hosts had significantly higher concentrations of alanine, gluconolactone, dopamine, putrescine, phenylalanine, glycerol, proline and quinic acid than oviparous aphids (ANOVA, df=1, p<0.05) (details of metabolite amounts measured from each morph are provided in Fig. S1). The second component of the PCA depicted the inter-individual variation of metabolites within each of the two morphs (Fig. 4).

**Figure 4:**
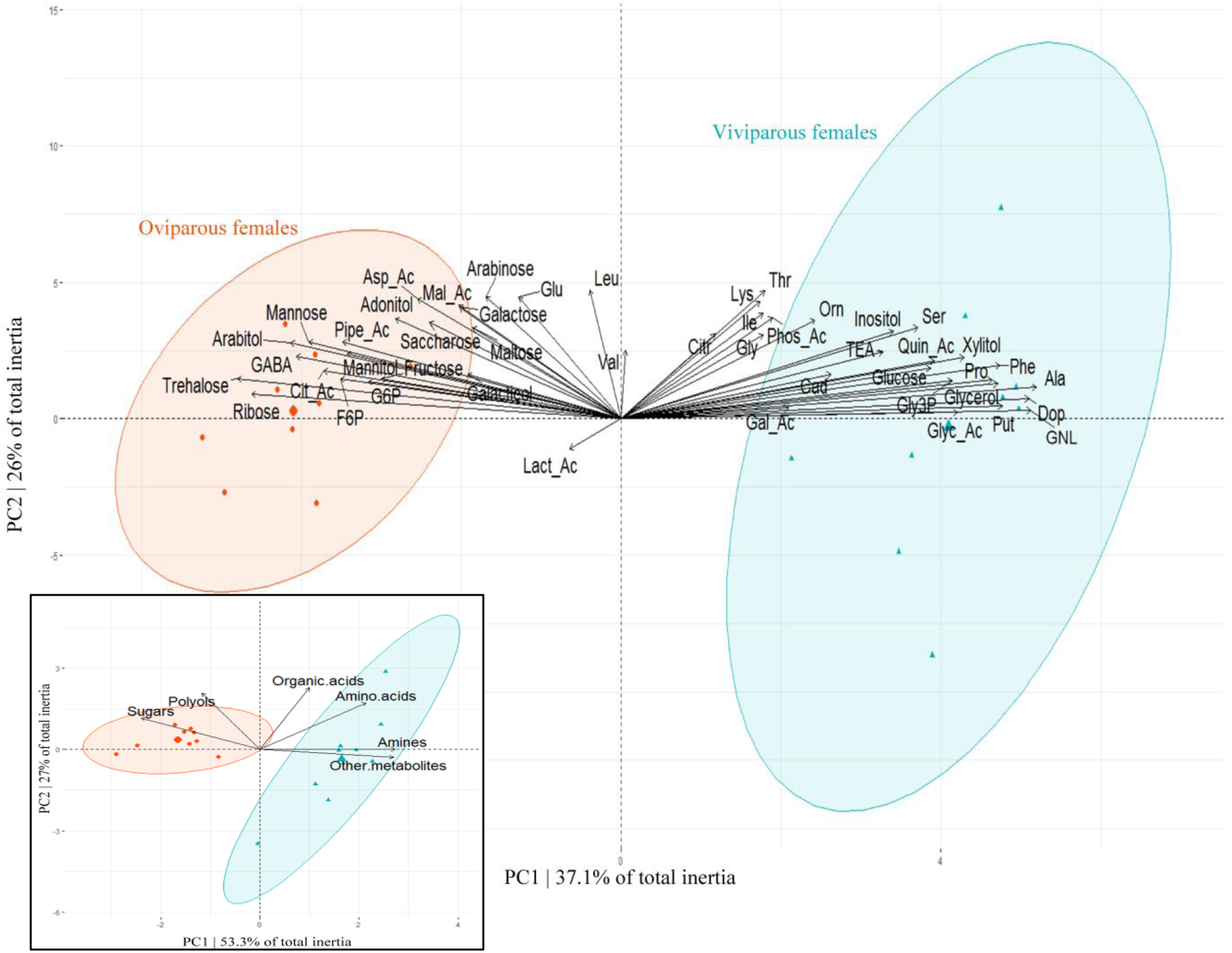
Multivariate analysis (PCA) on the first two principal components (PC) representing links between metabolic compounds (47 log-transformed variables, nmol.mg^-1^) and two aphid morphs (oviparous *vs*. viviparous females) of *Acyrthosiphon pisum*. Enclosed figure in the upper panel shows a PCA of the six metabolite categories. Confidence ellipses (95%) are constructed around each aphid group centroid (n=9 replicates by morph). Contributions of metabolite variables to PC1 and PC2 are provided in supplementary figure S3. Abbreviations are Alanine (Ala), Aspartic acid (Asp_Ac), Cadaverine (Cad), Citric acid (Cit_Ac), Citrulline (Citr), Dopamine (Dop), Fructose-6-phosphate (F6P), Galacturonic acid (Gal_Ac), Gamma aminobutyric acid (GABA), Gluconolactone (GNL), Glucose-6-phosphate (G6P), Glutamic acid (Glu), Glyceric acid (Glyc_Ac), Glycerol-3-phosphate (Gly3P), Glycine (Gly), Isoleucine (Ile), Lactic acid (Lact_Ac), Leucine (Leu), Lysine (Lys), Malic acid (Mal_Ac), Ornithine (Orn), Phenylalanine (Phe), Phosphoric acid (Phos_Ac), Pipecolic acid (Pipe_Ac), Proline (Pro), Putrescine (Put), Quinic acid (Quin_Ac), Serine (Ser), Threonine (Thr), Triethanolamine (TEA), Valine (Val).

The analysis by metabolic family revealed that sugars / sugar phosphates (at the exception of glucose) and polyols were measured in higher amounts in oviparous morphs, while amino acids, amines and other metabolites were generally found in higher concentrations in viviparous hosts (Fig. S2). Altogether, metabolic differences among oviparous and viviparous females revealed that activities of the pathways involved in aminoacyl-tRNA biosynthesis and glutathione metabolism were higher in viviparous females.

Oviparous hosts had a higher fat content ratio (mg fat/mg dry mass) than viviparous ones (0.63 ±0.02 and 0.51 ±0.03, n= 52 and n= 23, respectively) (ANOVA, LR=8.0, df=1, p<0.005). The fat mass represented 37.8 ±0.8% and 33.3 ±1.3% of the dry mass of oviparous and viviparous morphs, respectively.

## Discussion

Species interactions greatly contribute in shaping arthropods’ seasonal ecological strategies, because species needs to synchronize or unsynchronize their life cycle with interacting partners or antagonists. However, biotic-induced diapause signals are poorly studied. A few cases of predator-induced diapause have been documented in arthropods (Ślusarczyk 1995, Kroon et al. 2008), such as in *Daphnia magna* (Diplostraca: Daphniidae) in which the production of diapausing eggs is stimulated by predator exudates and chemicals originating from injured conspecifics (Ślusarczyk 1999). Reversely, low prey density was reported to influence summer diapause of the lady beetle *Hippodamia undecimnotata* (Coleoptera: Coccinellidae) (Iperti and Hodek 1974). Also, in herbivorous insects that require strong synchrony with their host plant phenology, resuming activities after winter diapause is also influenced by the physiological status of the plant (Leather et al. 1993). Similarly, the host plays a major role in parasitoid seasonal ecology. In addition to abiotic factors, such as photoperiod and temperature, the host genotype, species, size, life-stage and abundance can modulate parasitoid diapause (Tauber et al. 1986, Danks 1987).

We report that parasitoids can use host oviparous morph as a cue for diapause induction, with higher diapause incidence (up to 20%) expressed in *A. ervi* developing in oviparous *A. pisum* females compared to viviparous conspecifics. This pattern is likely due to differences in host physiology and metabolic contents. However, we have observed relatively high intrapopulation variability within each female’s offspring in response to the host morph, and to a lower extent in response to abiotic conditions, through the study of reaction norms. Polymorphism in the response to diapause-inducing cues (i.e., in plasticity) is known to be responsible for variability in diapause levels within populations experiencing different environmental conditions, but is still to be more deeply explored. As expected, parasitoids from the harsh winter environment expressed higher diapause levels than parasitoids from the mild winter environment. Of significance, only parasitoids from the harsh winter area and exposed to summer-like conditions relied on host morph as a cue for diapause induction.

Parasitoid populations of *A. ervi* from contrasted climatic environments (Canada and France) do not respond the same way to abiotic (photoperiod and temperature) and host cues. The French population of *Aphidius* spp. evolved under warming temperature conditions over the past decades, and this has allowed individuals of this species to remain active under mild winter conditions prevailing in this area, with none or small proportions of individuals entering diapause (Tougeron et al. 2017b). In mild winter areas, non-diapausing parasitoids maintain their populations by exploiting asexual anholocyclic aphid hosts during winter periods (Langer and Hance 2000, Andrade et al. 2015, 2016) as sexual morphs are rare in these areas (Dedryver et al. 2001). Diapause expression can be genetically lost or reduced in insects when they do not experience the necessary environmental factors for its induction (e.g., Bradshaw and Holzapfel 2001, Gariepy et al. 2015). Consequently, parasitoid populations from mild winter areas may not have evolved a response to sexual hosts, or they may have lost this capacity under changing environments.

The opposite pattern is observed in Canadian populations, where all aphid parasitoids enter diapause during winter (Brodeur and McNeil 1994). In these cold temperate regions, sexual morphs of aphids are produced at the end of the growing season, and represent the last hosts available for parasitoids before the onset of unfavorable winter conditions. In addition, parasitism of aphid sexual morphs on primary host plants allows parasitoids to overwinter nearby their hosts, thereby favoring host availability in spring for newly emerged parasitoids, and improving reproductive-cycles synchronization (Höller 1990, Christiansen-Weniger and Hardie 1997). In regions with harsh winter climates, parasitoids have coevolved with the seasonal occurrence of host morphs and may use oviparous morphs as a convergent signal with temperature and photoperiod decrease in fall to enter diapause. Canadian Aphidiinae parasitoids begin to overwinter as early as mid-July, with all individuals being in diapause by early September (Brodeur and McNeil 1994, Tougeron et al. 2018). This seasonal pattern might be an adaptation to avoid early lethal frosts. Moreover, we showed that oviparous hosts only influenced diapause under summer-like conditions, suggesting that encountering this morph informs the parasitoids for upcoming deleterious conditions and modulates diapause expression. In natural settings, alternative host species can be present, and both anholocyclic and holocyclic aphid populations can coexist (Dedryver et al. 2001), which may send confounding signals to parasitoids, and may explain why only a fraction of the population responded to oviparous morphs. In Canada, oviparous morphs of the pea aphid are present in the environment as soon as August (Lamb and Pointing 1972). In fall-like conditions, the morph effect was overridden by the temperature/photoperiod effect, which remains the main signal for diapause induction. Alternative diapause-inducing cues such as those associated with the host are usually viewed as factors modulating diapause expression, which is mainly triggered by temperature and photoperiod (Tauber et al. 1986). For example, in the polyphagous herbivore *Choristoneura rosaceana* (Lepidoptera: Tortricidae), diapause is dependent upon photoperiod and temperature, but under similar abiotic conditions, the proportion of larvae entering diapause differs depending on the host-plant species (Hunter and McNeil 1997). Moreover, the effect of the host-plant was observed even under photoperiod and temperature conditions known to induce low levels of diapause (Hunter and McNeil 1997). The relative importance of each environmental cue for diapause induction in insects remains to be evaluated for a significant number of species.

The response of parasitoids to host morph could be partly shaped by maternal effects, as females have the capacity to assess host quality through a combination of physiological, morphological, behavioural and chemical cues (van Baaren and Nénon 1996, Boivin et al. 2012). Developing immature parasitoids may also directly respond to the quality and quantity of metabolites available from hosts, which could trigger the onset of diapause. The overwintering metabolic and physiological syndrome is highly conserved among insects (Tauber et al. 1986), and both hosts and parasitoids may respond to the same molecules involved in diapause initiation. As an example concurring to this hypothesis, diapausing prepupae of the aphid parasitoid *Praon volucre* (Hymenoptera: Braconidae) showed similar proportions of some sugars (e.g. trehalose, fructose) and polyols (e.g. arabitol) (Colinet et al. 2012) than non-parasitized oviparous morphs of the pea aphid tested in our study. Our results suggest that high concentrations of some polyols and sugar metabolites in the oviparous morphs, as well as accumulation of fat reserves associated with the overwintering process, may either directly contribute to induce diapause in parasitoids developing in such hosts or may trigger the internal physiological cascade responsible for parasitoid diapause.

In the present work, oviparous *A. pisum* females have higher fat reserves than their viviparous counterparts. This finding is consistent with the metabolic phenotypes of the hosts, which revealed higher levels of sugar and sugar phosphate metabolites from the glycolytic pathway in oviparous females, this pathway providing elementary bricks for fatty acid and triacylglyceride (TAG) synthesis. Fatty acids serve as a main source of energy for physiological or ecological processes, including flight, gametes production, egg maturation and hormones synthesis (Arrese and Soulages 2010), and have been shown to represent up to 30% of aphids’ fresh mass (Dillwith et al. 1993, Sayah 2008). Interestingly, lipids can provide energy for overwintering insects and sugars can be metabolized to produce sugar-based cryoprotectant molecules (Storey and Storey 1991, Hahn and Denlinger 2011, Sinclair and Marshall 2018). In oviparous females, the need for TAG may be higher than in viviparous ones, as eggs with yolk (vitellus) are mostly composed of fat and proteins (Brough and Dixon 1990). Also, reserves from the fat-body, including TAG and glycogen, play major roles in overwintering insects, including diapause (reviewed in Sinclair and Marshall 2018) and could explain why oviparous aphids have high fat content to prepare their eggs for successful overwintering. Diapause entails important energetic costs for insects (Ellers and Van Alphen 2002, Hahn and Denlinger 2011) and they may enter diapause only when a critical body-mass or amount of energetic reserves has been reached (Colinet et al. 2010); for parasitoids, developing in an oviparous host could contribute to reach this level.

Metabolites acting as compatible solutes greatly contribute to insect cold hardiness and overwintering survival (Storey and Storey 1991, Bale 2002, Hodkova and Hodek 2004). Metabolic analyses identified sugars and polyols in higher amounts in oviparous females containing eggs intended to overwinter. Overwintering eggs of the aphid *Hyalopterus pruni* (Homoptera: Aphididae) are characterized by high values of mannitol and trehalose (Sömme 1969), as also observed in our *A. pisum* oviparous morphs. Glucose-6-phosphate and fructose were found at high concentrations in oviparous morphs of *A. pisum* and are precursors of sorbitol (Storey and Storey 1991), a cryoprotective compound also observed in diapausing individuals of *P. volucre* parasitoids (Colinet et al. 2012). Fructose-6-phosphate is a precursor of mannitol, and both are cryoprotectant molecules (Storey and Storey 1991) highly concentrated in oviparous female hosts, and found in most of overwintering insects (Leather et al. 1993). These metabolites may be responsible for diapause induction in parasitoids developing in oviparous morphs. Gamma aminobutyric acid was more concentrated in oviparous females and could also serve as an indirect seasonal cue for parasitoids because this neurotransmitter is known to be involved in insect perception of photoperiodic changes (Vieira et al. 2005).

Surprisingly, in viviparous females, we found high concentrations of glycerol, a cryoprotective compound usually associated with the diapause syndrome (Hayward et al. 2005). As suggested by the high concentrations of glucose observed in these females, glycogen production through gluconeogenesis pathway could be used as main source of energy by these viviparous morphs (Dixon 1985). In addition, observed physiological differences between host morphs are not necessarily linked to overwintering strategies. For example, viviparous aphids have high concentrations of proline, which is used as fuel for insect flight (Teulier et al. 2016). Viviparous aphids can rapidly produce winged individuals for dispersal in case of overcrowding or degradation of host plant quality (Hardie 1980).

To conclude, intra-and interspecific interactions are of primary importance for ecosystem functions, such as biological control, but still require deeper investigations in the context of diapause and seasonal strategies. Overwintering strategies are rapidly shifting in the context of climate change (Bradshaw and Holzapfel 2001, Bale and Hayward 2010) and may cause temporal mismatches between trophically interacting species (Tylianakis et al. 2008, Walther 2010). Thus, potential bottom-up effects on diapause, such as reported in our study, should be given more attention and should be considered as a potential factor explaining the low levels of diapause expression in insects from mild winter areas, together with global warming (Jeffs and Lewis 2013, Andrade et al. 2016, Tougeron et al. 2017b). In addition, there was variation for plasticity in diapause induction among female genotypes, mostly in response to the parasitized morph but also to abiotic conditions, as determined by slopes of the reaction norms. This means that there is genetic polymorphism in diapause plasticity within populations, which may allow natural selection to act in the context of rapid environmental and climate changes (Sgrò et al. 2016). Moreover, our results are of significance for the manipulation of insect diapause; e.g., in the context of mass rearing for the food industry, or for the biological control industry. More generally, a better appreciation of the processes governing phenology is needed to predict the consequences of such phenology changes on species interactions and synchrony across multiple trophic levels, community functioning and ecosystem services.

## Data accessibility and supplementary material

Metabolomics and diapause data have been made publicly available as a supplementary material attached to this publication. Supplementary figures S1, S2, S3 have also been made available: biorxiv.org/content/10.1101/371385v4.supplementary-material

## Supporting information

Public data

## Acknowledgements

We are grateful to both reviewers and both recommenders from PCI Ecology who made an excellent job in reviewing our manuscript and in providing strong advices on how to improve it. We thank G. Le Trionnaire at INRA Le Rheu for providing the aphid clones. We thank S. Llopis and J. Doyon for technical support and J. Jaquiéry for stimulating discussions. KT was funded by the Fyssen foundation, by the French Région Bretagne (ARED grant) and by the Canada Research Chair in Biological Control awarded to JB.

