## Supplementary material for "Sex makes them sleepy: host reproductive status induces diapause in a parasitoid population experiencing harsh winters": Public data: Data_PUBLIC_Readme.txt

EXCEL SHEET #1

*********************
KEY FOR TABLE READING
*********************

	pop		CANADA or FRANCE				the origin of the parasitoid population
	batch		1 or 2						temporal replication, as described in the method section of the mansucript
	female		1 to 12						parasitoid female ID
	morph		Oviparous, Control or Viviparous		reproduction status of the aphid morph
	condition	17 �C | L10:D14 or 20 �C | L16:D8		abiotic conditions to which parasitized aphids were exposed
	mummies		number from 1 to 8				number of aphid mummies produced by each parasitoid female at each condition
	diapause	number from 0 to 8				number of diapausing parasitoids within the mummies

----------------------------------------------------------------------------------------------------------------------------------------------------

EXCEL SHEET #2

	group	O or V		indicates the group (Oviparous or Viviparous) to which each aphid was attributed
	ID	A to I		individual identification of the tested aphid


********************************
LIST OF METABOLITE ABBREVIATIONS
********************************


Alanine (Ala), 
Aspartic acid (Asp_Ac), 
Cadaverine (Cad), 
Citric acid (Cit_Ac), 
Citrulline (Citr), 
Dopamine (Dop), 
Fructose-6-phosphate (F6P), 
Galacturonic acid (Gal_Ac), 
Gamma aminobutyric acid (GABA), 
Gluconolactone (GLN), 
Glucose-6-phosphate (G6P), 
Glutamic acid (Glu), 
Glyceric acid (Glyc_Ac), 
Glycerol-3-phosphate (Gly3P), 
Glycine (Gly), 
Isoleucine (Ile), 
Lactic acid (Lact_Ac), 
Leucine (Leu), 
Lysine (Lys), 
Malic acid (Mal_Ac), 
Ornithine (Orn), 
Phenylalanine (Phe), 
Phosphoric acid (Phos_Ac), 
Pipecolic acid (Pipe_Ac), 
Proline (Pro), 
Putrescine (Put), 
Quinic acid (Quin_Ac), 
Saccharose, 
Serine (Ser), 
Threonine (Thr), 
Triethanolamine (TEA), 
Valine (Val).


----------------------------------------------------------------------------------------------------------------------------------------------------
