## Supplementary material S1, S2 and S3 for "Sex makes them sleepy: host reproductive status induces diapause in a parasitoid population experiencing harsh winters"

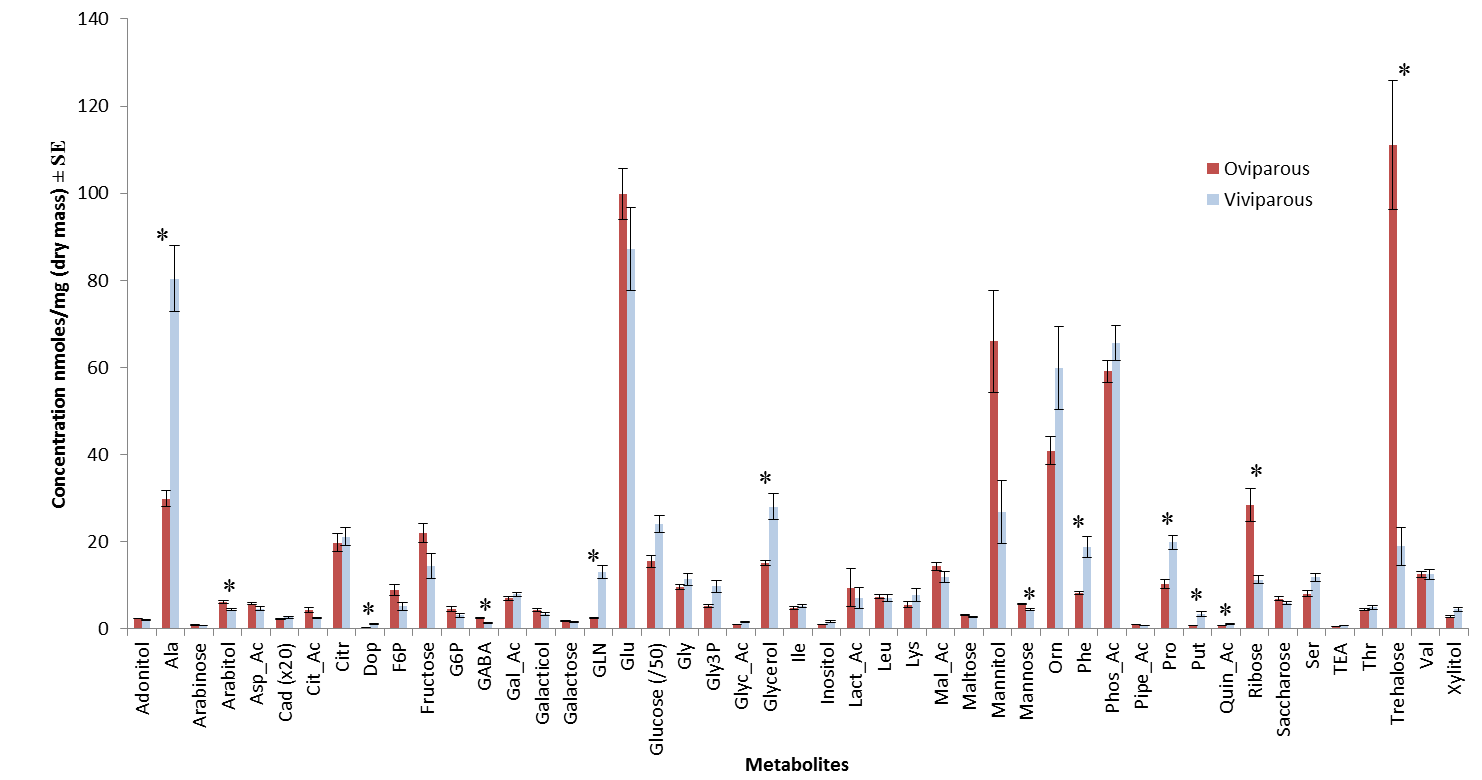

Figure S1: Mean concentration (nmoles.mg^-1^) ±SE of each of the 47 metabolite observed by gas-chromatography spectrometry in oviparous females (red) and viviparous females (blue) of the pea aphid *Acyrthosiphon pisum*. Note that cadaverine levels (Cad) have been multiplied by 20 and glucose levels have been divided by 50 on this graph to improve scaling. Alanine (Ala), Aspartic acid (Asp_Ac), Cadaverine (Cad), Citric acid (Cit_Ac), Citrulline (Citr), Dopamine (Dop), Fructose-6-phosphate (F6P), Galacturonic acid (Gal_Ac), Gamma aminobutyric acid (GABA), Gluconolactone (GNL), Glucose-6-phosphate (G6P), Glutamic acid (Glu), Glyceric acid (Glyc_Ac), Glycerol-3-phosphate (Gly3P), Glycine (Gly), Isoleucine (Ile), Lactic acid (Lact_Ac), Leucine (Leu), Lysine (Lys), Malic acid (Mal_Ac), Ornithine (Orn), Phenylalanine (Phe), Phosphoric acid (Phos_Ac), Pipecolic acid (Pipe_Ac), Proline (Pro), Putrescine (Put), Quinic acid (Quin_Ac), Serine (Ser), Threonine (Thr), Triethanolamine (TEA), Valine (Val). N=9 replicates/morph. Stars (*) indicate significant differences between aphid morphs (p<0.05, ANOVA with FDR-adjusted p-values) for each metabolite.

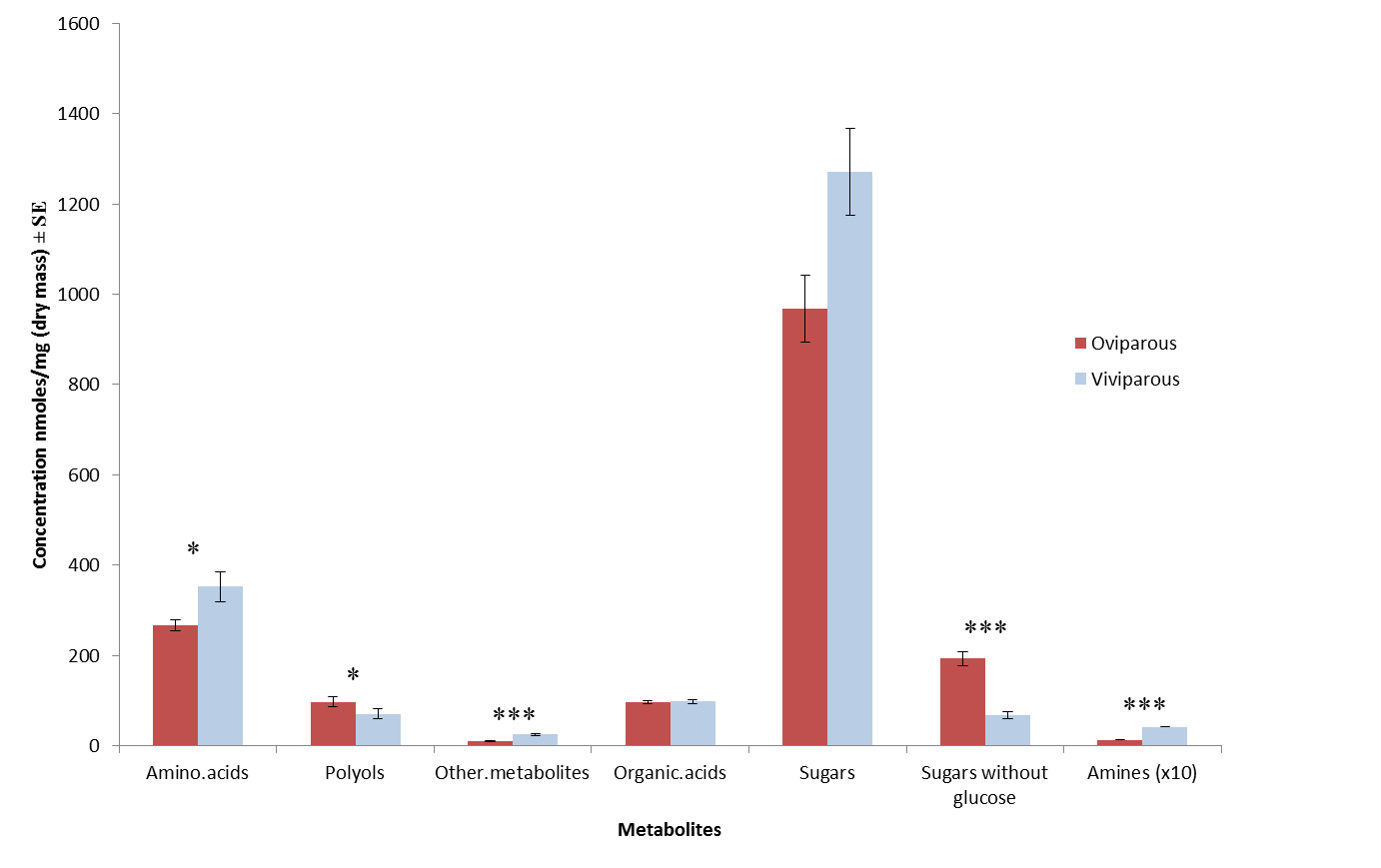

Figure S2: Mean concentration (nmoles.mg^-1^) ±SE of each of the 7 metabolite groups observed by gas-chromatography spectrometry in oviparous females (red) and viviparous females (blue) of the pea aphid *Acyrthosiphon pisum*. Note that amines levels have been multiplied by 10 to improve scaling. Sugars are represented with and without glucose because this metabolite has an order of magnitude more than ten times superior to other sugars. N=9 replicates/morph. Stars indicate significant differences between aphid morphs (*** p<0.001, * p<0.1, ANOVA with FDR-adjusted p-values) for each metabolite group.

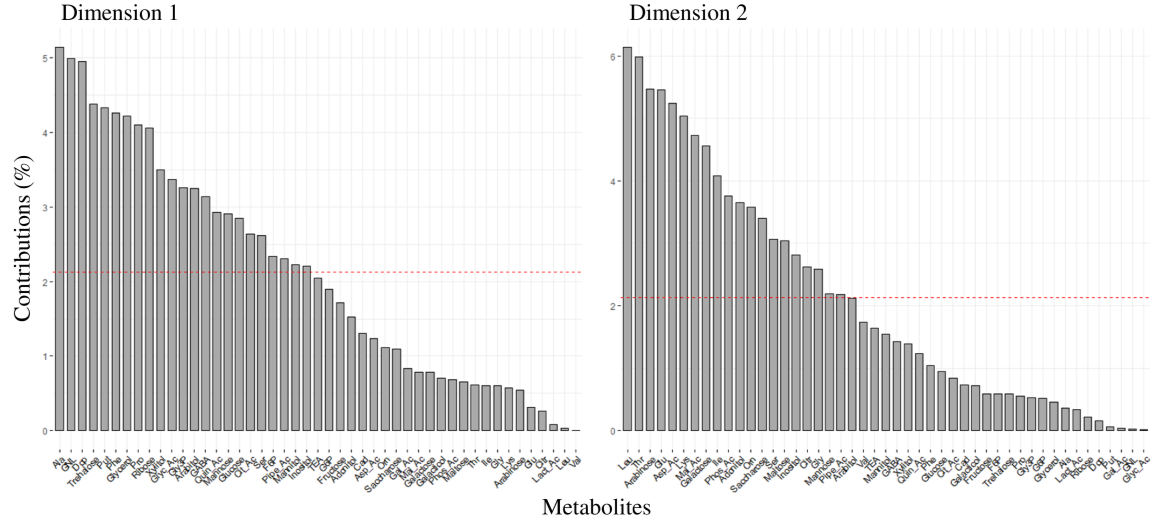

Figure S3: Contributions of metabolite variables to PC1 and PC2 of the PCA presented in Figure 3. Abbreviations are as follow: Alanine (Ala), Aspartic acid (Asp_Ac), Cadaverine (Cad), Citric acid (Cit_Ac), Citrulline (Citr), Dopamine (Dop), Fructose-6-phosphate (F6P), Galacturonic acid (Gal_Ac), Gamma aminobutyric acid (GABA), Gluconolactone (GNL), Glucose-6-phosphate (G6P), Glutamic acid (Glu), Glyceric acid (Glyc_Ac), Glycerol-3-phosphate (Gly3P), Glycine (Gly), Isoleucine (Ile), Lactic acid (Lact_Ac), Leucine (Leu), Lysine (Lys), Malic acid (Mal_Ac), Ornithine (Orn), Phenylalanine (Phe), Phosphoric acid (Phos_Ac), Pipecolic acid (Pipe_Ac), Proline (Pro), Putrescine (Put), Quinic acid (Quin_Ac), Serine (Ser), Threonine (Thr), Triethanolamine (TEA), Valine (Val).
